# Daily Heat Stress Induces Accumulation of Non-functional PSII-LHCII and Donor-side Limitation of PSI via Downregulation of the Cyt *b*_6_*f* Complex

**DOI:** 10.1101/2025.11.06.687104

**Authors:** Laura Laihonen, Tutta Tomberg, Linda Vuorijoki, Lauri Nikkanen, Paula Mulo, Marjaana Rantala

## Abstract

The rapidly warming climate is driving increasingly frequent and intense heat waves worldwide, posing major challenges to plant metabolism and growth. Despite the central role of photosynthesis in plant growth, the effects of prolonged heat stress on the primary reactions of photosynthesis remain poorly understood. Here, we studied how long-term heat exposure affects growth and photosynthetic performance in the model plant *Arabidopsis thaliana*. Plants were exposed daily to high temperature (38°C) for four hours throughout the growth period. PSII-LHCII complexes remained structurally intact, but ΔF/F_m_’, electron transfer rate (ETRII) and proportion of open reaction centers (qL) were lower under prolonged heat. Strikingly, the content of the Cytochrome *b*_6_*f* complex, which mediates electron transfer between photosystems, was reduced by 30–40%. Consequently, donor-side limitation of PSI was higher and the overall PSI electron transfer rate (ETRI) was lower under long-term heat. Further, carbon assimilation was significantly lower under long-term heat exposure, but PSI acceptor side limitation was not affected. We propose that downregulation of the cytochrome *b*_6_*f* complex restricts electron delivery to PSI, thereby preventing PSI over-reduction due to reduced carbon assimilation.

**Summary statement:** Heat-acclimated *Arabidopsis thaliana* plants accumulate non-functional PSII-LHCII supercomplexes. PSI complex is protected against over-reduction by restricting electron flow from Cyt *b_6_f* as an acclimatory response.

## Introduction

The global mean temperature in 2024 was 1.5°C above the pre-industrial era (WMO, 2025). Global warming has led to more frequent and severe heatwaves, disrupting ecosystems and threatening agriculture. Based on different modelling and field-warming experiments, it was estimated that each degree-Celsius increase in global mean temperature would reduce global yields of wheat by 6.0%, rice by 3.2%, maize by 7.4%, and soybean by 3.1% (Zhao et al., 2017). The rapidly warming climate, particularly the increased frequency and intensity of heatwaves (IPCC, 2021), significantly impacts plant metabolism. Understanding the mechanisms of heat damage on plant metabolism is fundamental for mitigation strategies that prevent the loss of plant productivity under increasing temperatures.

Photosynthesis is considered to be among the most prone cell functions to be damaged upon heat stress (Allakhverdiev et al., 2008). Light-driven photosynthetic electron transfer is conducted by membrane-embedded large protein complexes: photosystem (PS) II and PSI, which contain peripheral light harvesting antennae (LHCII and LHCI, respectively), and are interconnected by cytochrome (Cyt) *b_6_f* and two mobile electron carriers: plastoquinone (PQ) and plastocyanin (PC). The electrons, originating from water, are transferred from one complex to another and ultimately used for reducing NADP^+^ to NADPH. Water oxidation by PSII and PQH_2_ oxidation by Cyt *b_6_f* releases protons in the lumen. The formed proton gradient across the thylakoid membrane is released by ATP synthase and ATP is formed. Generated NADPH and ATP are used in downstream metabolism, most importantly to assimilate atmospheric CO_2_ in the Calvin-Benson-Bassham (CBB) cycle. Balancing the energy supply from electron transfer reactions with the downstream metabolic demands is essential for efficient photosynthesis under fluctuating environmental conditions. To cope with environmental fluctuations, plants have evolved mechanisms to regulate their photosynthetic apparatus according to environmental cues. These mechanisms involve adjustments in the light harvesting antennae to balance the amount of excitation energy between PSII and PSI (state transitions), dissipation of excess absorbed energy as heat (NPQ) and regulation of the electron transfer chain at Cyt *b_6_f* and PSI (photosynthetic control and cyclic electron transfer).

Excessive heat affects all aspects of plant photosynthesis from electron transport to carbon metabolism (Berry and Björkman, 1980; Inoué et al., 1987; Sharkey, 2005; Sharkey and Zhang, 2010). Particularly carbon assimilation is reduced due to impaired activity of Rubisco activase (RCA) (Feller et al., 1998; Law and Crafts-Brandner, 1999; Crafts-Brandner and Salvucci, 2000; Rokka et al., 2001). The effects of high temperature stress on photosynthetic light reactions are less understood. The reported effects of high temperature on light reactions include dissociation of the peripheral LHCII antenna from PSII core (Armond et al., 1980; Sundby et al. 1986; Srivastava et al., 1997; Mathur et al., 2011), destacking of thylakoid membranes (Gounaris et al., 1983; 1984), more oxidized PQ-pool (Pshybytko et al., 2008) and impairment of oxygen evolving complex (OEC) (Santarius, 1975; Enami et al., 1994). Indeed, PSII has long been considered to be the most prone ETC component to high temperature stress (Berry and Björkman, 1980), in comparison with the thermostable PSI (Ivanov et al., 2017). Notably, many of the existing studies are based on acute single exposure *in vitro* experiments (Gounaris et al., 1983; 1984; Sundby et al., 1986; Inoué et al., 1987; Enami et al., 1994; Semenova, 2004 Komayama et al., 2007), while understanding of how long-term, recurring heat exposure experienced by whole plant []such as that during a heatwave[] impacts photosynthesis *in vivo* remains limited.

Here, we studied the effects of physiologically relevant long-term heat stress on plant growth and photosynthetic reactions. *Arabidopsis thaliana* plants were subjected daily to 38°C for four hours at mid-day during their whole life cycle. In this setup, plants begin to acclimate to long-term recurring elevation in temperature, but also experience short-term stress due to daily heat periods. The effects of daily heat exposure on plant growth, electron transfer reactions and photosynthetic proteins and protein complexes were studied using chlorophyll-fluorescence and absorbance-based methods, immunoprobing and native-PAGE. We show that under these conditions both photosystems remain structurally intact, but their photochemistry is impaired. Intriguingly, we show that daily heat stress results in dramatic downregulation of the Cyt *b_6_f* complex and thus donor-side limitation of PSI. Moreover, heat-acclimated plants showed significantly reduced carbon assimilation rates. We propose that in heat-acclimated plants the adjustment of the stoichiometry of ETC components acts as a protective mechanism, restricting electron flow from PSII to protect PSI complex from overreduction, with a concomitant reduction of photosynthetic activity.

## Materials and methods

### Plant growth conditions and phenotyping

*Arabidopsis thaliana* ecotype Columbia-0 (Wt) was grown under 12 h light/ 12 h dark cycle at photosynthetic photon flux density (PPFD) of 130 μmol m^-2^ s^-1^ (VALOYA LUMI-CS LL120 LED), + 23°C and relative humidity of 50% for 4-5 weeks. Heat-acclimated plants were grown in a chamber where temperature elevated daily from 23 to 38°C for four hours and RH was 50-70% depending on the temperature. Light intensity and cycle were same as in Ctrl.

The phenotypes of heat-acclimated and Ctrl plants were imaged weekly throughout the growth period. The images were processed with Adobe Photoshop (version 26.6.1, Adobe Systems Inc.).

### Pigment analysis (DMF)

Leaf discs (63.6 mm^2^) collected from 5-week-old Wt plants grown under daily heat stress and Ctrl conditions were immersed in *N,N*-dimethylformamide (DMF), then incubated overnight in darkness. Leaf debris was pelleted by centrifugation (5 min, 20 000 x g, RT), and absorbances of the supernatants were measured at 663.8, 646.8, and 480 nm with spectrophotometer (UV-1800, Shimadzu). Pigment concentrations were calculated according to Wellburn (1994) using the following equations: Chl *a* = 12A663.8 - 3.11A646.8, Chl *b* = 20.78A646.8 – 4.88A663.8, Car = (1000A480 – 1.12Chl *a* – 34.07Chl *b*)/245. Nine leaf discs from three biological replicates were measured (n=27).

### Transmission electron microscopy (TEM)

Leaf samples for TEM were prepared according to Hyman & Jarvis (2011). Briefly, the leaf fragments (1 x 1 x 5 mm) from 5-week-old wild-type *Arabidopsis thaliana* plants were cut at 90 ° angle to the midrib from leaves of same developmental stage. The samples were collected in the middle of light and heat period. Ten leaf fragments from three individual plants were dissected in fixative solution (2.5% (w/v) glutaraldehyde, 0.1 M sodium phosphate buffer, 4.0% (w/v) paraformaldehyde, pH 7.2) and incubated for four hours at room temperature. Samples were then transferred to 4 °C and stored overnight. The sample preparation for TEM was done according to Hyman & Jarvis (2011). Samples were analyzed with JEM-1400 Plus Transmission Electron Microscope.

### Thylakoid isolation

Thylakoid proteins were isolated according to Järvi et al., 2011. Chl concentration was determined from the samples according to Porra et al., 1989. Isolations were done at the middle of heat period.

### Polyacrylamide gel electrophoresis (PAGE)

For SDS-PAGE, thylakoid proteins were solubilized with Laemmli buffer (Laemmli, 1970) supplemented with 6 M urea and 10% β-mercaptoethanol and run on 12% acrylamide gels. The proteins were electroblotted onto Immobilon-FL membrane (Merck Millipore) with Trans-blot® (Bio-Rad) device, blocked with 5% milk or bovine serum albumin (BSA) in TTBS (20□mM Tris–HCl, pHC7.5, 150□mM NaCl, and 0.05% Tween 20) and incubated with primary antibody overnight. The sample was loaded as follows: 0.5□μg of Chl for Phospho-Thr/Ser antibody (Cell Signaling Technology, 9381S; 1:6000) and 1 μg of Chl / 10 μg of protein for D1 (DE-loop antibody (Kettunen et al., 1996); 1:8000), PsbO-1 (AS142824, 1:5000), PSAB (AS10695; 1:2500, Agrisera), ATPB (AS05085; 1:5000, Agrisera), Cyt f (AS06119; 1:1000, Agrisera), Rieske (AS08330; 1:5000) and Lhcb1 (AS01004; 1:2000), NdhL (2 µg Chl, gift from T.Shikanai, Shimizu et al., 2008, 1:5000) Rubisco L (AS03037, 1:5000) Rubisco activase (gift from T.J. Andrews, Rokka et al., 2001, 1:1000). Samples for most of the SDS–PAGE analyses were normalized to chlorophyll content, as this approach is commonly used to account for differences in thylakoid membrane protein abundance. Since heat grown plants exhibited lower chlorophyll content, we repeated subset of analyses with protein-based loading, and yielded broadly comparable results (Fig S1). The membrane was incubated with a secondary antibody IRDye® 800CW Goat anti-Rabbit IgG (1:20 000 in 1% milk/TTBS) and scanned with Li-Cor Odyssey CLx.

For BN-PAGE, gels and samples were prepared as in Rantala et al., 2018. Thylakoid samples were solubilized with 1% digitonin or β-DMM, and samples containing 5 µg of Chl were separated on BN gels.

### Chlorophyll a fluorescence and P700 absorbance spectroscopy

DKN spectrophotometer (Heinz Walz GmbH) was used to measure Chl *a* fluorescence from PSII with pulse-modulated 540 nm measuring light, and redox changes of P700, PC and Fd were measured by monitoring absorbance changes at 780–820, 820–870, 840–965 and 870–965 nm. PC, P700 and Fd redox changes were deconvoluted based on differential model plots (DMPs) (Klughammer and Schreiber, 2016) measured specifically for Arabidopsis leaves. Normalization of PC, P700 and Fd redox changes was done according to the maximal redox changes that were determined with the software’s NIRMAX script. Steady state oxidation and reduction levels were calculated as relative PC_ox_, P700_ox_ and Fd_red_. Light response curves were measured by illuminating leaves with 635 nm red actinic light for 5 min with the following light intensities: 100, 200, 500, 750 and 1200 μmol m^−2^ s^−1^. Plants were dark-adapted for 1 h prior measurement, and measurements on heat-acclimated plants were done in the middle of heat treatment period. Minimal and maximum fluorescence (F_o_ and F_m_) and the maximum P700 signal were determined in the dark-adapted state. The effective PSII quantum yield of illuminated samples (ΔF/F_m_’) was calculated as (F_m_’ - F)/F_m_’ (Genty et al. 1989), the fraction of open PS II centers (qL) was calculated as qL = (F_m_’ - F) × F_o_/((F_m_ - F_o_)/F_m_ + (F_o_/F ^’^))/(F_m_’ - (F_o_/((F_m_ - F_o_)/F_m_ + (F_o_/F ^’^)) × F (Oxborough and Baker 1997; Kramer et al., 2004), relative electron transfer rate of PSII as ETR (II) = (ΔF/F_m_’) × PPFD × 0.84 × 0.5 (Genty et al., 1989), and non-photochemical quenching as NPQ = (F_m_ - F_m’_)/F_m_’ (Demmig-Adams, 1990).

### 77K measurement

77K fluorescence measurements were performed on thylakoids diluted in BTH buffer (25 mM BisTris/HCl (pH 7.0), 20% (w/v) glycerol and 0.25 mg·ml^1^ Pefabloc) to a chlorophyll concentration of 5 µg/ml. Thylakoids kept in liquid nitrogen and were excited with blue (440 nm) light and fluorescence emission was detected with QEPro spectrophotometer (Ocean Optics). Spectra were normalized to 685 nm peak.

### ECS measurements

The ECS signal arising from light-induced shifts in the absorbance spectra of thylakoid pigments was measured by simultaneously monitoring the absorbance differences at 546 nm and 520 nm with a JTS-150 pump-probe spectrophotometer (BioLogic). The detectors were shielded form scattering effects by BG-39 filters (Schott). Actinic light and single-turnover flashes were provided by a 660 nm 10W laser diode (AeroDiode). The ECS signal was deconvoluted from scattering artefacts as (A_520 nm_ x 1.611) - (A_546 nm_ x 1.247) according to the scaling factors determined by Kramer and Sacksteder (1998). ECS_T_, light-induced ECS amplitude corresponding to the magnitude of light-induced *pmf*, was calculated as the difference between ECS in light (130 μmol m^−2^ s^−^1) and an Y_0_ value obtained from the first-order exponential fit to the decay kinetics of the ECS signal during 350 ms dark intervals. Dark intervals were given every 15 s. The ECS signal was normalized to the ΔECS induced by a 20 µs saturating single-turnover flash administered prior to the onset of actinic illumination. The gH^+^ parameter (thylakoid membrane conductivity to protons) was calculated as the inverse of the time constant of the first-order exponential fit to ECS decay kinetics during a dark interval (Avenson et al., 2005; Cruz, 2005). The vH^+^ parameter (the rate of proton flux over the thylakoid membrane) corresponding to the initial slope of the decay of the ECS signal upon cessation of illumination was calculated as *pmf* x gH^+^ (Cruz, 2005). Partitioning of *pmf* between its component parts ΔpH and ΔΨ was determined by monitoring the post-illumination kinetics of the ECS signal, and by quantifying the magnitude of the rapid decay of the signal below the dark baseline, caused by efflux of protons through ATP synthase, and the gradual increase in the signal in the dark caused by influx of counter-ions (Cruz, 2005). Plants were dark-adapted for 1 h prior measurement and measurements on heat-acclimated plants were done both at the middle of heat treatment period and before heat treatment period.

### Cytochrome f redox change measurements

Cytochrome *f* redox kinetics were measured as time-resolved absorbance changes at 554 nm with a 546 and 573 nm baseline (ΔA_554 nm_ – ((ΔA_546 nm_+ ΔA_573 nm_)/2) during the same light regime used for the ECS measurements, with the JTS-150 spectrophotometer (BioLogic). BG39 filters (Schott) were used to shield the detectors from scattering effects. Cytochrome *f* re-reduction rates were determined by fitting the post-illumination decay kinetics of the signal to a first-order exponential function y = y_0_ + A_1_ × exp(-(x - x_0_)/t_1_), with *k*_Cyt_ *_f_* _red_ = 1/t_1_. Maximal oxidation of Cyt *f*, determined by the initial decrease in the deconvoluted signal upon illumination, was used to normalise the Cyt *f* oxidation values.

### Gas exchange measurements

Carbon assimilation measurements were performed using 5-week-old plants grown at a PPFD of 130 µmol m^−2^ s^−1^ with the GFS-3000 gas exchange system (Walz) equipped with 3,010□M standard measuring head and 3,041□L LED light source. Measurements were performed on 6 plants/condition at a humidity of 14,000 ppm, at CO_2_ concentration of 400 ppm, and the cuvette temperature was set to 20°C. CO_2_ assimilation and H_2_O evaporation were monitored at three different light intensities (200, 600 and 2000□µmol m^−2^ s^−1^) for 5 min/intensity.

## Results

### Long-term daily heat exposure results in stunted growth and reduced content of photosynthetic pigments

To study the effects of daily heat exposures on photosynthesis, plants were grown in chamber where temperature was elevated daily for four hours from 23°C to 38°C throughout their lifecycle. Control (Ctrl) plants were grown at constant 23°C. The plants were photographed weekly from week one to five (Fig. 1a) and the biophysical and biochemical characterization was conducted on plants at the end of the growth period, at week five. As demonstrated in Fig. 1a (lower panel), daily heat treatment resulted in stunted growth, fresh weight of the plants reaching only ∼40% of that of Ctrl plants (Fig. 1b), as well as altered rosette morphology with elongated petioles and leaves elevated from the soil. Additionally, strong hyponastic leaf movement was observed during the heat exposures: the petioles started to bend upwards from the soil during the high temperature period (Supplementary video). Further, in heat-acclimated plants, the accumulation of chlorophyll (Chl) *a* and *b* was reduced by ∼30% with no significant effects on Chl *a/b* ratio, as compared to Ctrl (Fig. 1c, 1d). Moreover, the accumulation of carotenoids (xantophylls and carotenes) was reduced by∼30% (Fig. S2).

**Figure 1.**
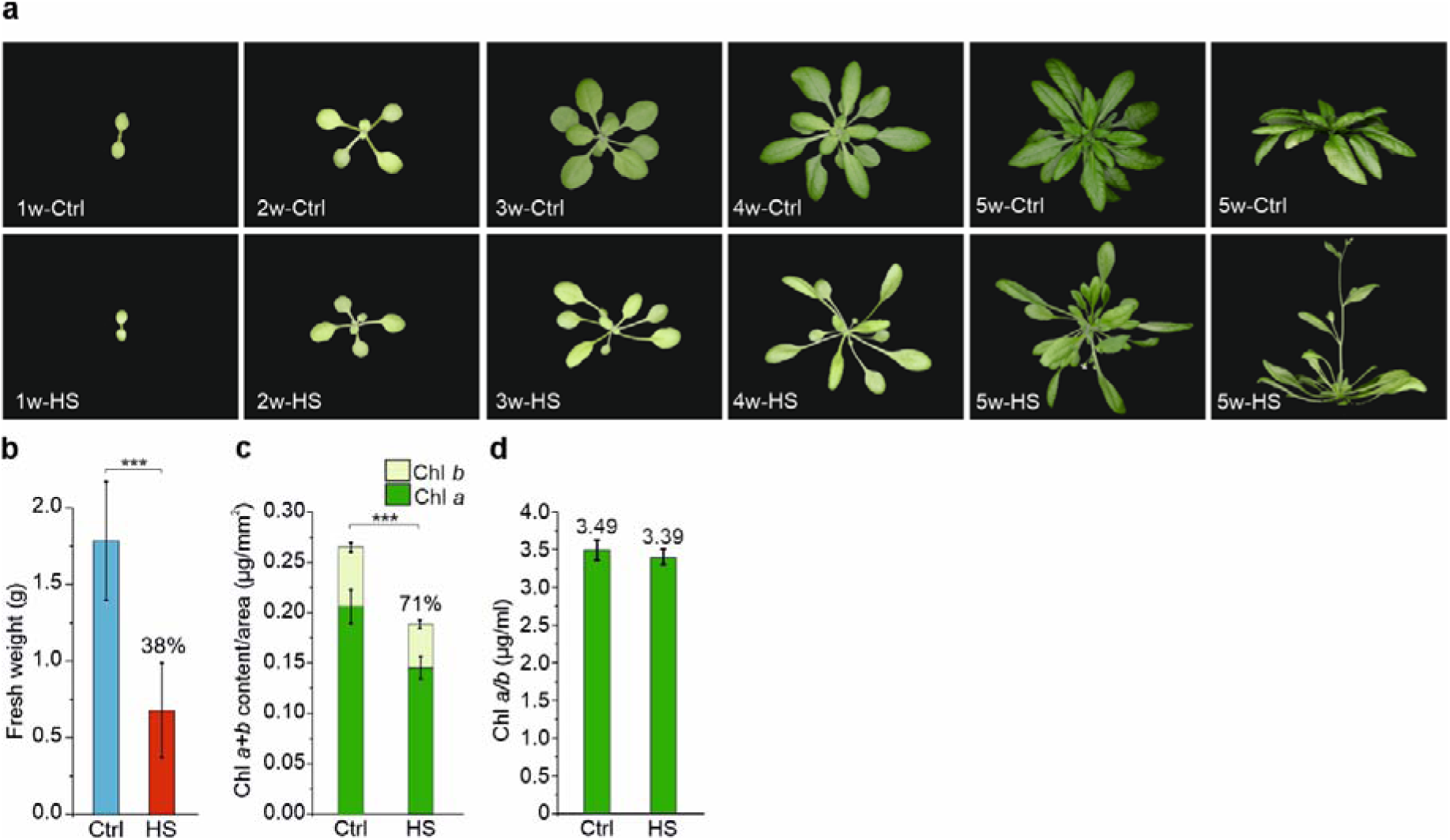
The effect of long-term daily heat exposure on (a) plant growth phenotype, (b) fresh weight (c) chlorophyll content and (d) chlorophyll a/b ratio. Plants were grown for five weeks in growth chamber where temperature increased daily from 23°C to 38°C for four hours (HS), whereas control (Ctrl) plants were grown at constant 23°C. Fresh weight (n = 15) and chlorophyll (Chl) content (n = 27) were measured at week five. Statistical significance according to Student’s t-test is indicated with an asterisk (* = < 0.05, ** = < 0.01, *** = <0.001). Standard deviations are indicated with error bars.

### Daily heat stress alters thylakoid ultrastructure and number of grana per chloroplast

Earlier reports have demonstrated complete linearization of the thylakoid membrane system upon sudden exposure to high temperature (Gounaris et al., 1983; 1984). Transmission electron microscopy was used to study the effects of daily heat exposure on the thylakoid membrane ultrastructure. Leaf samples were collected from 5-week-old control plants and from plants grown under long-term heat stress in the middle of the heat period. As demonstrated in Figure 2, heat-acclimated plants showed higher number of grana per chloroplast (52 ± 2) compared to Ctrl conditions (33 ± 10), but the average height of grana was significantly smaller in heat-acclimated plants (107.6 ± 60 nm) compared to Ctrl plants (120.4 ± 60 nm) (Fig. 2c). Additionally, daily heat exposure resulted in larger plastoglobuli as compared to Ctrl conditions (Fig. 2a, b).

**Figure 2.**
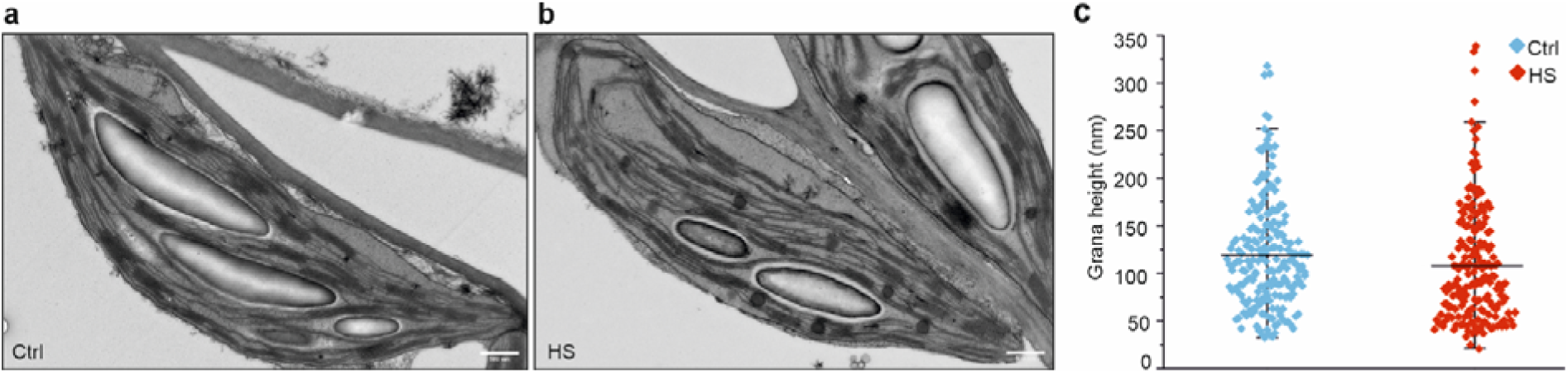
Transmission electron microscopy analysis of chloroplast ultrastructure. a, b) Representative electron micrographs of chloroplasts from control (Ctrl) and heat-acclimated (HS) plants. 8000 × magnification was used; 500 nm scale bar is indicated at bottom-right corner. c) Height of individual grana stacks was measured from 202 (Ctrl) and 194 (HS) grana stacks. Samples were collected in the middle of light and heat period. Mann-Whitney Rank Sum Test showed statistical significance between Ctrl and HS plant grana height (p = 0.014). Standard deviations are indicated with error bars.

### PSII and PSI are structurally intact but functionally impaired under prolonged heat stress

Previous studies have demonstrated that PSII-LHCII complex is the most prone ETC component to high temperature-induced damage. Heat stress has been shown to result in LHCII disconnection from PSII (Armond et al., 1978; 1980) and degradation of the PSII core protein D1 as well as disassembly of the OEC complex (Enami et al., 1994; Yamashita et al 2008). To investigate the effects of daily moderate heat exposures on PSII-LHCII supercomplexes, thylakoid membranes from heat-acclimated and Ctrl plants were isolated and solubilized with non-ionic detergent β-dodecyl maltoside (β-DDM), followed by separation with Blue Native (BN)-PAGE. In contrast to earlier studies, heat-acclimated plants showed enhanced association of LHCII to PSII core, represented by high accumulation of large forms of PSII-LHCII supercomplexes, namely C2S2M(2), containing PSII core dimer (C2) attached to two strongly (S2) and one (M) or two (M2) moderately bound LHCII trimers (Fig. 3a). Moreover, heat-acclimated plants accumulated reduced levels of detached M-LHCII complexes and PSII monomers (Fig. 3a). However, no differences were observed in the overall abundance of the PSII reaction center protein D1 or the LHCII protein Lhcb1 between heat-acclimated and Ctrl plants (Fig. 3b). Notably, the PSII core protein CP43 as well as LHCII proteins were less phosphorylated in heat-acclimated plants than in Ctrl (Fig 3b).

**Figure 3.**
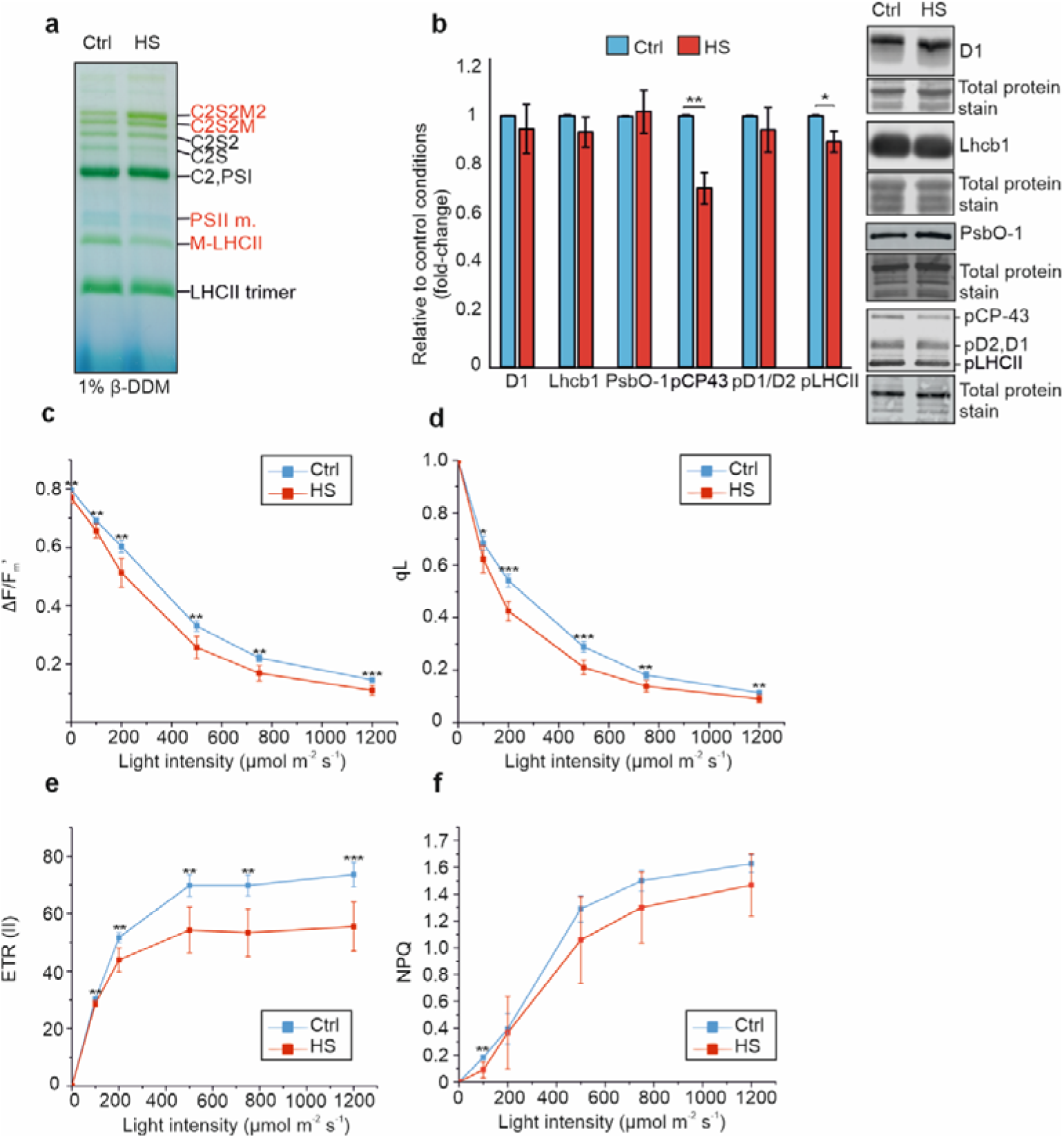
The effects of daily heat stress (HS) on PSII-LHCII complex. a) Representative BN gel for thylakoids isolated from control (Ctrl) and HS plants using β-dodecyl maltoside (β-DDM) for solubilization. 5 µg of Chl was loaded per lane. Complexes showing differences between Ctrl and HS are marked in red. b) Western blots and protein quantification from thylakoid membrane proteins isolated from Ctrl and HS plants using antibodies against D1, Lhcb1 and Phosphorylated Thr/Ser. Quantifications are normalized to Ctrl. (n = 3 biological replicates) The membranes were stained with total protein stain to verify equal loading. c) – f) Rapid light response curves were recorded using chlorophyll *a* fluorescence with Dual KLAS-NIR system. ΔF/F_m_’, proportion of open PSII reaction centers (qL), PSII electron transport rate (ETR) and non-photochemical quenching (NPQ) were measured. Statistical significance according to Student’s t-test is indicated with an asterisk (* = < 0.05, ** = < 0.01, *** = <0.001). (n = 6 individual plants). Experiments (thylakoid membrane isolation and DKN measurements) were done at mid-day in the middle of heat period. Standard deviations are indicated with error bars.

Despite accumulating structurally intact C2S2M2 supercomplexes, heat-acclimated plants showed higher initial fluorescence (F_o_) and reduced maximal fluorescence (F_m_) in chlorophyll *a* fluorescence measurement (Supplemental Data S1). Elevated F_o_ under heat stress has been reported previously (e.g., Havaux 1993; Pospíšil et al. 1998) and suggests a blockage of PSII. Havaux (1993) proposed that the PSII donor side is affected already at 32°C, whereas inhibition of electron transfer from Q_A_ to Q_B_ occurs only at 42°C or higher temperatures. However, at the protein level, we did not observe differences in the accumulation of the OEC subunit PsbO (Fig. 3b). Chlorophyll *a* fluorescence measurement in the light-adapted state showed significantly lower ΔF/F_m_′ (in which F_m_′ indicates fluorescence during light saturating point and F indicates fluorescence before the saturating flash), electron transport rate through PSII (ETR(II)), and the proportion of open PSII reaction centers (qL) in heat-acclimated plants compared to controls (Fig. 3c-e). Since no differences were observed in the abundance of PSII core proteins or physical disconnection of the LHCII antenna from the PSII core (Fig 3a, b), it seems that a fraction of PSII-LHCII complexes is inactive, but not degraded. NPQ was not markedly affected in heat-acclimated plants (Fig. 3f) indicating reduced photochemistry of PSII. Further research needed to characterize the mechanism of PSII damage.

PSI is considered to be relatively stable under acute heat stress (Ivanov et al., 2017), and previous studies have demonstrated enhanced LHCII phosphorylation and increased absorbance cross-section of PSI in darkness (Ivanov and Velitchkova, 1990; Mohanty et al., 2002; Nellaepalli et al., 2011). To study PSI composition, thylakoid membranes isolated from heat-acclimated and Ctrl plants were solubilized with digitonin in ACA buffer, which preserves the labile interaction of LHCII with PSI, while solubilizing complexes from the grana stacks, grana margins and stroma lamellae (Rantala et al., 2017). In BN gels (Fig 4a), no differences were observed in the amount of PSI complex, but Western blot analysis indicated higher abundance (∼20%) of PSI reaction center subunit PsaB in the heat-acclimated plants (Fig. 4b). Although there were no defects in the PSI accumulation, PSI quantum yield (Y(I)) and electron flow through PSI (ETR(I)) were lower in heat-acclimated plants at nearly all light intensities (Fig. 4c, d). Moreover, in line with the reduced LHCII phosphorylation level (Fig. 3b), heat-acclimated plants exhibited slightly lower accumulation of the state transition -specific PSI-LHCII (Fig. 4a). To study the relative excitation of PSII and PSI, low temperature fluorescence spectra was recorded. Interestingly, increased F735(PSI)/F685(PSII) ratio was observed in heat acclimated plants (Fig. 4e), indicating higher relative excitation of PSI.

**Figure 4.**
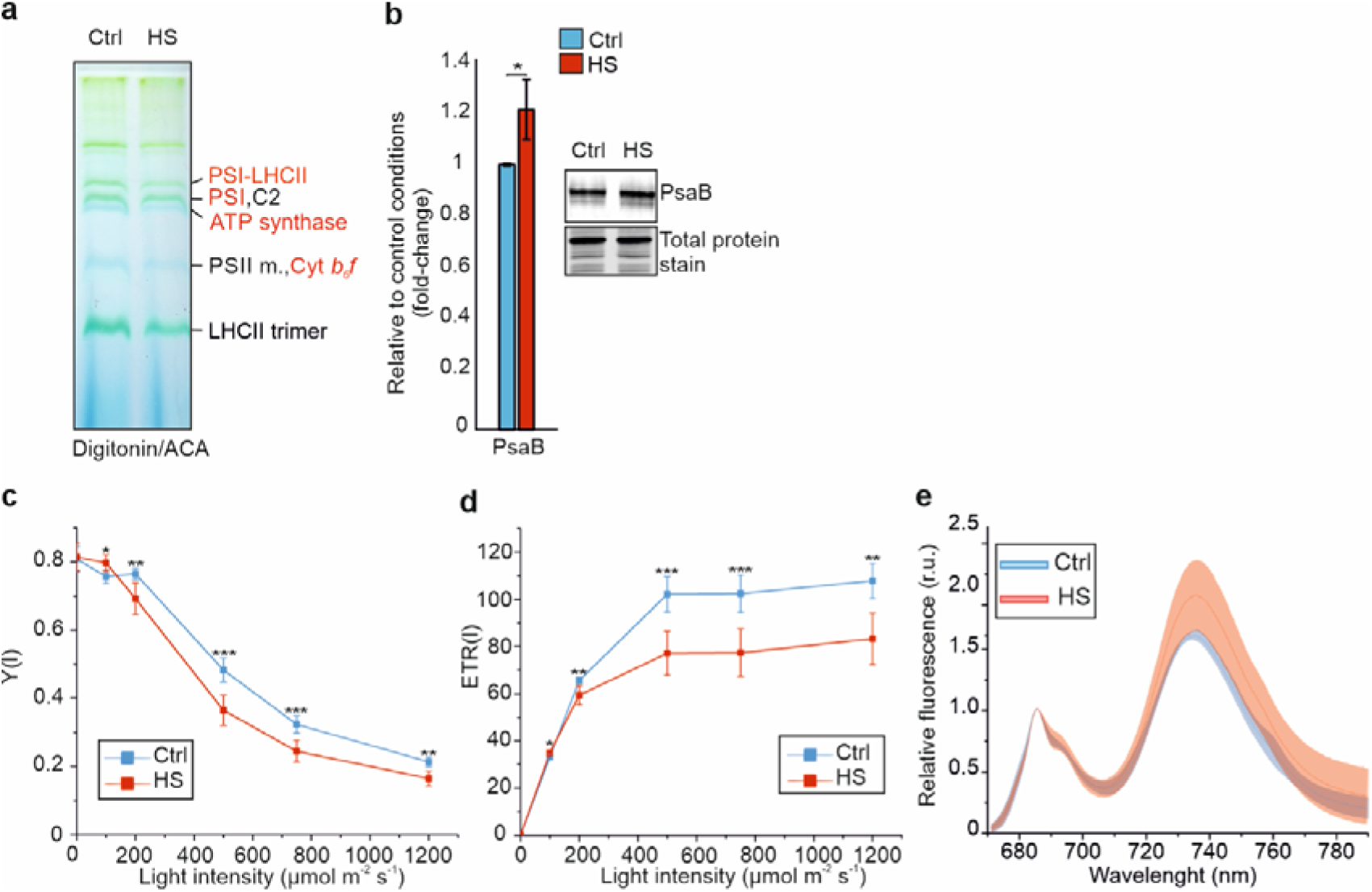
The effects of daily heat stress (HS) on photosystem PSI. a) BN-PAGE analysis of native protein complexes from thylakoids of control (Ctrl) and heat acclimated (HS) plants using digitonin in ACA buffer for solubilization. 5 µg of Chl was loaded per lane. Complexes showing differences between Ctrl and HS are marked in red. b) Western blot analysis and quantification of PsaB protein from thylakoids isolated from Ctrl and HS. The quantifications were normalized to Ctrl. (n = 3 biological replicates). c) – d) Rapid light response curves were recorded using P700 absorption spectroscopy with Dual KLAS-NIR system to record. PSI quantum yield (Y(I)) and PSI electron transport rate (ETR) were recorded (n = 6 individual plants). e) 77K fluorescence spectra from Ctrl and HS thylakoids (n = 3 biological replicates). Statistical significance according to Student’s t-test is indicated with an asterisk (* = < 0.05, ** = < 0.01, *** = < 0.001). Experiments (thylakoid membrane isolation and DKN measurements) were done at mid-day in the middle of heat period. Standard deviations are indicated with error bars.

### Downregulation of Cyt b_6_f complex limits electron transfer to PSI donor side

To determine whether the declined PSI photochemistry detected upon daily heat exposures (Fig. 4) resulted from defects in PSI donor or acceptor side, we measured the acceptor and donor side limitations of PSI and the relative redox states of P700, PC and Fd (Fig. 5 a – c). Figure 5a shows that Y(NA) across all light intensities is lower in heat-acclimated plants than in Ctrl, indicating no limitations at PSI acceptor side. This result is in line with the less reduced Fd in heat-acclimated plants as compared to Ctrl plants (Fig. 5c). In contrast, donor-side limitation, Y(ND), was substantially increased in heat-acclimated plants (Fig. 5b). Moreover, both PC and P700 were more oxidized (Fig. 5c), implying that the electron flow from Cyt *b_6_f* to PSI is restricted, resulting in PSI donor side limitation. This is indeed supported by the dramatically reduced Cyt *b_6_f* content in heat-acclimated plants, as shown by BN-PAGE (Fig. 4a). This result was confirmed by 30-40% decrease in the content of CytF and Rieske subunits of the Cyt *b_6_f* complex in Western blot analysis (Fig. 5d). In addition, the ATP synthase complex (Fig. 4a) and the ATP synthase subunit ATPβ (Fig. 5d) were significantly less abundant in heat-acclimated plants as compared to the Ctrl.

**Figure 5.**
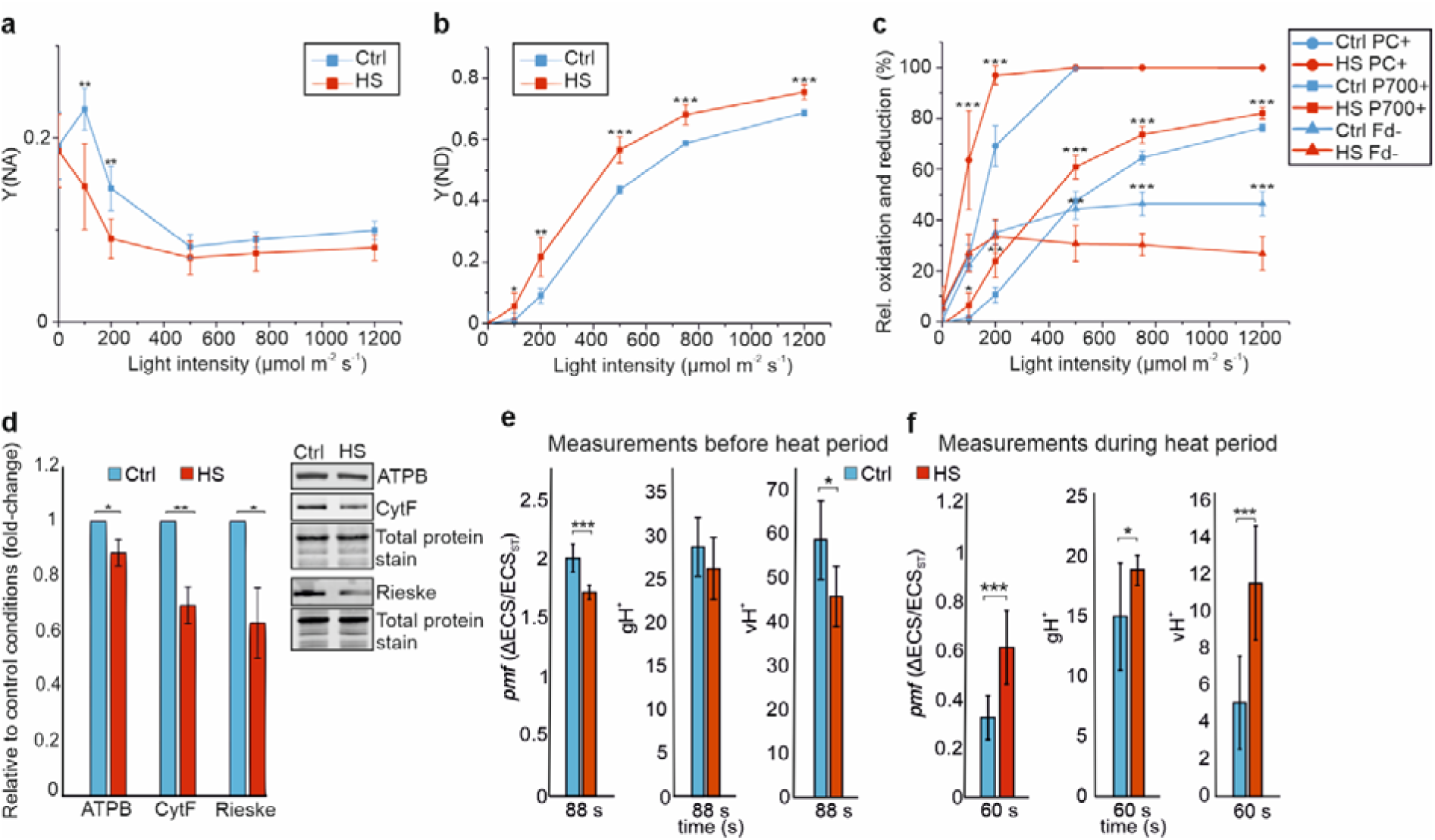
Heat stress (HS) effects on donor and acceptor side limitation of PSI and proton motive force (*pmf*). a) – c) Dual KLAS-NIR was used to measure PSI acceptor side limitation (Y(NA), donor side limitation (Y(ND) and relative oxidation/reduction of plastocyanin (PC), P700 and ferredoxin (Fd). (n = 6 individual plants). d) Western blots and protein quantifications from thylakoids of control (Ctrl) and HS plants using antibodies against ATPB, Cyt F and Rieske. (n = 3 biological replicates). Thylakoid membrane isolation and DKN measurements were conducted at mid-day in the middle of heat period. e) – f) ECS was measured during transitions from dark to 130 μmol photons m^-2^ s^-1^ of green actinic light. Absorbance difference between 546 and 520 nm (electrochromic shift, ECS) was measured with a JTS-150 spectrometer, and light-induced *pmf* was determined from the dark interval relaxation kinetics of the ECS signal. Conductivity of the thylakoid membrane (gH^+^) after 88 s or 60 s of illumination was determined as the inverse of the time constant of first-order post-illumination decay kinetics of the ECS signal. Thylakoid proton flux (vH^+^), calculated as *pm*f × gH^+^. ECS measurements were done both before the heat period and during the heat period to distinguish acute short-term effects of heat treatment from long-term acclimatory responses. Statistical significance according to Student’s t-test is indicated with an asterisk (* = < 0.05, ** = < 0.01, *** = <0.001). (n = 6 individual plants). Standard deviations are indicated with error bars.

To analyze the effects of downregulated Cyt *b_6_f* and ATP synthase on proton motive force (*pmf*), Cyt f redox kinetics and electrochromic shift (ECS), respectively, were monitored during dark to light transition (Fig. 5; Fig. S3; Fig. S4). In order to distinguish short-term heat-induced differences from the long-term acclimatory responses resulting from decreased Cyt *b_6_f* levels, measurements were done before the onset of heat period (Fig. 5e; Fig. S3a - d) as well as at mid-day in the middle of the heat period (Fig. 5f; Fig. S3e - g). In concordance with the reduced Cyt *b_6_f* levels, heat-acclimated plants showed decreased *pmf* compared to Ctrl plants (Fig. 5e; Fig. S3a) when measurements were conducted before the onset of heat period. Additionally, thylakoid membrane conductivity (gH^+^) and proton flux to the lumen (vH^+^) were calculated from ECS measurements between 15 s intervals. There were no major differences in gH^+^ between Ctrl and HS plants, but vH^+^ was significantly reduced in HS plants (Fig. 5e; Fig. S3c, d) in agreement with the decreased PSII and PSI electron transport rates. In contrast to measurements made before heat period, heat-acclimated plants showed higher *pmf,* and significantly higher vH^+^ compared to Ctrl plants when measurements were conducted during heat period (Fig. 5f; Fig. S3e – g). This finding may indicate enhanced CET during the heat period. This hypothesis was supported by immunoblot analysis showing increased amount of NdhL, a subunit of the CET-related PSI-NDH complex (Peng and Shikanai, 2011), in the heat-acclimated plants compared to Ctrl plants (Fig. S5).

During the heat period, a slightly higher proportion of Cyt *f* remained oxidized during dark-to-light transitions in the heat-acclimated plants in comparison to control plants (Fig. S4a), while the reduction rates of Cyt *f* were similar between treatments (Fig. S4b). These results suggest increased electron flux through Cyt *b*_6_*f* during the heat period in the heat-acclimated plants, consistent with the hypothesis of increased CET PSI.

### Heat-acclimated plants show reduced carbon assimilation rates

Since high temperatures have been shown to inhibit photosynthesis by decreasing Rubisco activity via inhibiting RCA (Crafts-Brandner and Salvucci, 2000; Salvucci et al., 2001), we assessed carbon assimilation capacity of heat-acclimated plants (Fig. 6a). Carbon assimilation rates as well as stomatal conductance were significantly reduced in heat-acclimated plants compared to Ctrl plants (Fig. 6). Interestingly, there were now significant differences in total RCA levels between Ctrl and HS plants, but in accordance with the previous findings (Rokka et al., 2001), more RCA was associated with the thylakoid membranes in HS plants (Fig. S6). No differences were observed in the total amount of Rubisco or in the amount of thylakoid membrane-associated Rubisco between Ctrl and HS plants (Fig. S6).

**Figure 6.**
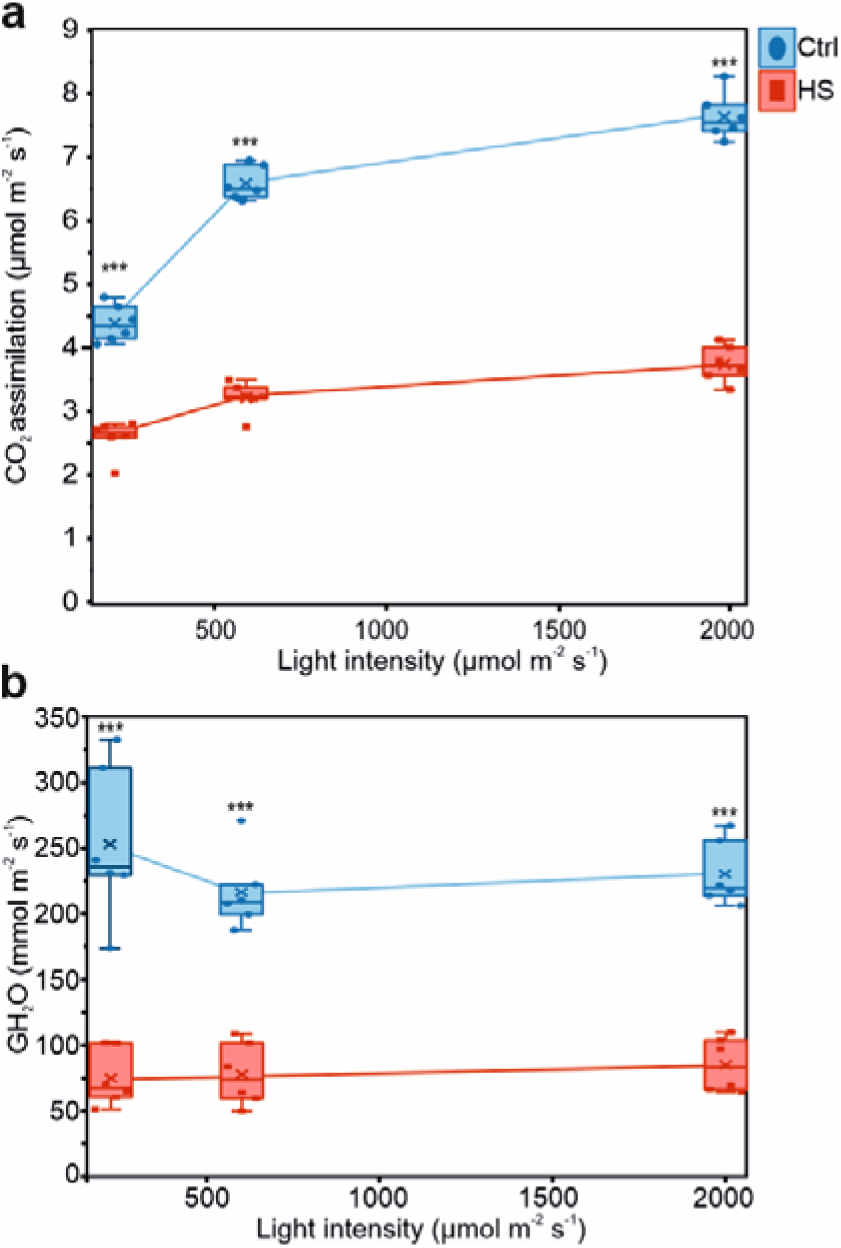
The effect of daily heat stress (HS) on carbon assimilation and stomata conductivity. Gas exchange measurements were performed with GFS-3000 (Walz). Measurements were done at mid-day using three different light intensities (200, 600 and 2000 μmol m^-2^ s^-1^) and illuminating leaf for 5 min with each intensity at 20°C. Statistical significance according to Student’s t-test is indicated with an asterisk (* = < 0.05, ** = < 0.01, *** = < 0.001). (n = 6 individual plants). Standard deviations are indicated with error bars.

## Discussion

Increase in ambient temperature affects multiple aspects of plant metabolism, especially photosynthetic reactions (Berry and Björkman, 1980). Currently, holistic understanding of the effects of physiologically relevant long-term heat stress on the thylakoid membrane reactions is completely lacking. Long-term changes in environmental conditions trigger changes in the consumption of ATP and NADPH, and the stoichiometry of photosynthetic complexes is dynamically regulated to meet the metabolic needs of the plant (Schöttler and Tóth, 2014). It has been suggested that these dynamic changes are mediated by the redox state of photosynthetic electron transfer chain, i.e. PQ pool and Cyt *b_6_f*, which act as a signaling network to regulate gene expression levels in nucleus and chloroplast (Anderson et al., 1997; Pfannschmidt, 2003).

Long-term daily heat stress leads to a pronounced reduction in chlorophyll (Chl) content (Fig. 1d), consistent with previous findings (Fatma et al., 2021; Mustafa et al., 2021). This decrease has been attributed to enhanced accumulation of chlorophyllase and Chl-degrading peroxidases (Rossi et al., 2017). Notably, the abundance of Chl-binding photosynthetic complexes, i.e. PSII–LHCII and PSI, remains unchanged (Fig. 3a, b; Fig. 4a, b), indicating that under long-term recurrent heat stress, fewer Chl molecules are associated with these complexes compared with Ctrl conditions. In heat-acclimated plants, the overall abundance of D1 and Lhcb1 remain unaltered. Strikingly, in plants grown under long-term heat stress, the C2S2M2 and C2S2M supercomplexes are more frequent forms of PSII-LHCII than in Ctrl plants, indicating enhanced association of LHCII with the structurally intact PSII core. This is in contrast with earlier reports demonstrating D1 degradation (Komayama et al., 2007; Yamashita et al., 2008) and physical disconnection of the LHCII antenna from the PSII core upon heat stress (Armond et al., 1980; Sundby et al. 1986; Srivastava et al., 1997; Mathur et al., 2011). Despite the presence of structurally intact supercomplexes, under elevated temperature, lower fraction of the captured light energy is utilized in photochemistry (lower ΔF/F_m_’ and ETRII), and correspondingly higher fraction is passively dissipated due to closed reaction centers (Fig. 3a-d). The accumulation of non-functional PSII–LHCII complexes in heat-acclimated plants may reflect reduced efficiency of the PSII repair cycle. Light-induced damage of the PSII core protein D1 results in D1 degradation followed by biosynthesis and co-translational insertion of the new D1 copy to the disassembled PSII complex. Specifically, damaged and phosphorylated PSII dimer monomerizes in the grana stacks and the monomers migrate to non-appressed thylakoid membranes. The PSII core proteins D1, D2 and CP43 are dephosphorylated (Koivuniemi et al., 1995; Rintamäki et al., 1996; Rokka et al., 2000) before the proteolytic degradation of the D1 protein. Next, the newly synthesized D1 protein is inserted into the PSII subcomplex and the new functional complex is assembled and translocated to the appressed grana. Finally, the LHCII trimers are associated with the PSII centers. In heat-acclimated plants, the PSII repair cycle intermediate, PSII monomer (Aro et al 2005), is almost completely absent (Fig. 3a), which supports the hypothesis of impaired PSII repair cycle.

Figure 4 (a,b) shows that daily heat stress does not affect the accumulation of the PSI complex and, in fact, increases the PSI reaction center protein PsaB content. However, association of PSI with LHCII antenna trimer is reduced (Fig. 4a), yet more excitation energy is directed to PSI (Fig. 4e) in heat-acclimated plants. Such increase of relative PSI excitation could indicate randomization of the thylakoid protein complexes and consequent energy spillover from PSII to PSI (Barber, 1980; Ivanov and Velitchkova, 1990). Our analysis of the thylakoid ultrastructure shows smaller grana stacks (height) in heat-acclimated plants, but not full linearization of the thylakoid membrane (Fig. 2), and therefore randomization of PSII and PSI is unlikely. More detailed analysis of the protein complex distribution in heat-acclimated plants is needed to understand the lateral heterogeneity of the complexes under heat stress conditions and the possible excitation energy spill-over from PSII to PSI.

Strikingly, we show that electron flow beyond Q_A_ is suppressed by dramatic Cyt *b_6_f* downregulation (Fig. 4a; Fig. 5d), which restricts electron flow to PSI and more oxidized P700. Decrease in the Cyt *b_6_f* content likely serves to protect PSI from excessive reduction and ROS generation, especially when carbon assimilation is significantly reduced (Fig. 6) and PSI cannot donate electrons to downstream electron acceptors. Protection of PSI is crucial, as it lacks similar repair machinery as PSII: once PSI is damaged, the repair process requires *de novo* synthesis and assembly of the entire complex (Sonoike, 2011). In agreement with the lower Cyt *b_6_f* content, we observed lower *pmf* in heat-acclimated plants compared to Ctrl plants (Fig. 5e) when measurements were conducted for heat-acclimated plants before the daily heat treatment. Despite the lower ATP synthase content (Fig. 4a; Fig. 5d), gH^+^ levels were similar between Ctrl and heat-acclimated plants (Fig. 5e; Fig. S3c). However, Rott et al. (2011) showed that ATP synthase content can be decreased down to 50 % without affecting gH^+^. Intriguingly, when *pmf* was measured during the heat period (Fig. 5f), proton flux to the lumen was higher in heat-acclimated plants (Fig. S3g). This might indicate increased CET that is activated during the heat period. Indeed, previous studies have reported activation of CET under high temperature (Bukhov et al., 1999; Schrader et al., 2004; Wang et al., 2006; Zhang and Sharkey, 2009; Zhang et al., 2023), which has been hypothesized to maintain sufficient ATP content upon heat-induced thylakoid leakiness. Alongside increased proton flux, increased electron flux through Cyt *b*_6_*f* and high accumulation of NdhL protein, representing CET-related PSI-NDH (Peng and Shikanai, 2011), were detected in heat-acclimated plants compared to Ctrl (Fig. 5f; Fig. S4a; Fig. S5). The results indicate that in addition to acclimatory responses resulting in decreased Cyt *b_6_f* levels, heat-acclimated plants might respond to daily increase in temperature via short-term regulatory mechanisms, such as CET.

Overall, our results demonstrate that the mechanisms limiting photosynthesis under single heat exposure differ from those activated upon long-term repeated exposures to 38°C, applied in our experimental setup. The long-term responses of the photosynthetic apparatus, include (i) decreased grana height, (ii) impaired function of PSII and PSI, despite their structural integrity, (iii) accumulation of non-functional PSII-LHCII complexes, likely due to defects in PSII repair cycle, (iv) reduced LHCII phosphorylation and association with PSI, (v) donor side limitation of PSI, and (vi) downregulation of Cyt *b_6_f*, which may serve as an acclimatory strategy to protect PSI under repeated heat exposures (Fig. 7).

**Figure 7.**
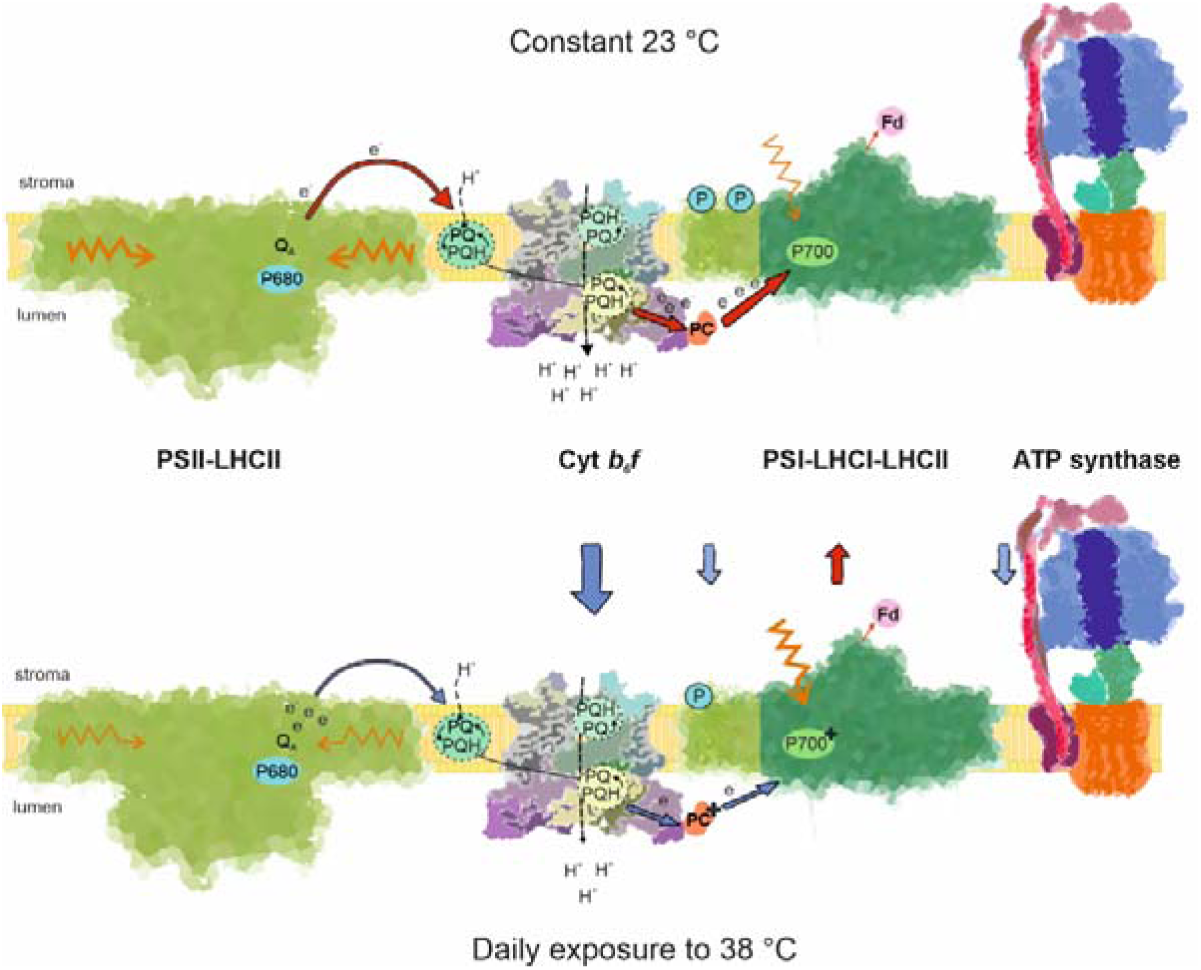
Summary of the effects of long-term daily heat exposure to photosynthetic light reactions. Compared to control conditions (constant 23°C), plants exposed daily to 38°C for four hours, exhibit pronounced changes in the structure and function of the photosynthetic machinery. Blue arrows indicate reduced (Cyt *b_6_f*, ATP synthase and state transition specific PSI-LHCI-LHCII complex), whereas red arrow indicate increased (PSI core subunit PsaB) accumulation of thylakoid protein complexes. Despite retaining structurally intact PSII–LHCII supercomplexes, heat-acclimated plants show reduced PSII photochemical efficiency. In addition, relative PSI excitation is elevated (orange arrows) despite the lower abundance of PSI–LHCI–LHCII complexes. Downregulation of Cyt *b_6_f* decreases electron flux to PSI, leading to more oxidized plastocyanin (PC) and P700.

## Acknowledgements

We thank Prof. Eva-Mari Aro for valuable advice on TEM sample preparation and for expert guidance with the imaging. Dr. Steffen Grebe, Dr. Laura Wey, Dr. Tapio Lempiäinen, Dr. Minna Konert, Dr. Esa Tyystjärvi and Prof. Gyozo Garab are thanked for discussions and valuable insights. Versa Elements is thanked for building the heat chamber and Mika Keränen, Tapio Ronkainen and Samuli Pyytövaara are thanked for the technical assistance. TEM specimen preparation and imaging was done at Laboratory of Electron Microscopy, the Institute of Biomedicine, University of Turku. Special thanks to Jenni Laine and Markus Peurla in technical guidance at TEM Laboratory. The PHOTOSYN infrastructure (University of Turku; https://sites.utu.fi/photosyn/) is acknowledged for the excellent research environment.

