## Supplementary Figures S1-S6 for "Daily Heat Stress Induces Accumulation of Non-functional PSII-LHCII and Donor-side Limitation of PSI via Downregulation of the Cyt *b*_6_*f* Complex"

### Laihonen et al., 2026

### Supplementary data and figures

### As separate files:

**Supplementary data S1.** $F_o$ and $F_m$ values measured from dark-acclimated control (Ctrl) and heat-acclimated (HS) plants. Values were measured from individual plants from 3 biological replicates (n = 16). Student's t-test was used to analyse statistical significance, and p-values are included in the file.

**Supplementary video 1.** Plant movement was monitored during the daily heat period. Timestamp at the upper-left corner represents time after onset of heat period, where temperature started to increase to 38°C where it remained for four hours until declining back to 23°C.

Fig. S1

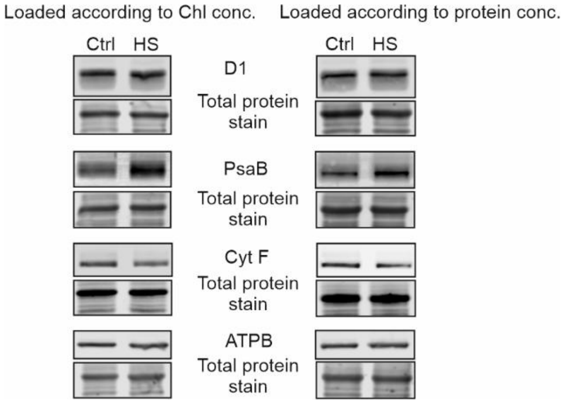


**Supplementary Figure 1.** To ensure that chlorophyll (Chl) concentration -based loading gives similar results as protein concentration -based loading despite decreased Chl content of heat-acclimated (HS) plants, thylakoid membrane proteins were run with SDS-PAGE and samples were loaded according to protein and Chl concentrations, and immunoprobed with D1, PsaB, CytF and ATPB antibodies. 1 µg Chl or 10 µg of protein was loaded/well.

Fig. S2

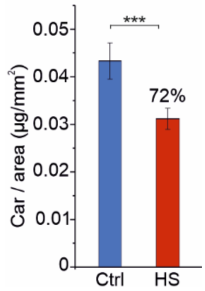


**Supplementary Figure 2.** Plants were grown for five weeks in growth chamber where temperature increased daily from 23 °C to 38 °C for four hours (HS), whereas control (Ctrl) plants were grown at constant 23 °C. Carotenoid content was measured at week five (n = 27). Statistical significance according to Student's t-test is indicated with an asterisk (* = < 0.05, ** = < 0.01). (n = 27). Standard deviations are indicated with error bars.

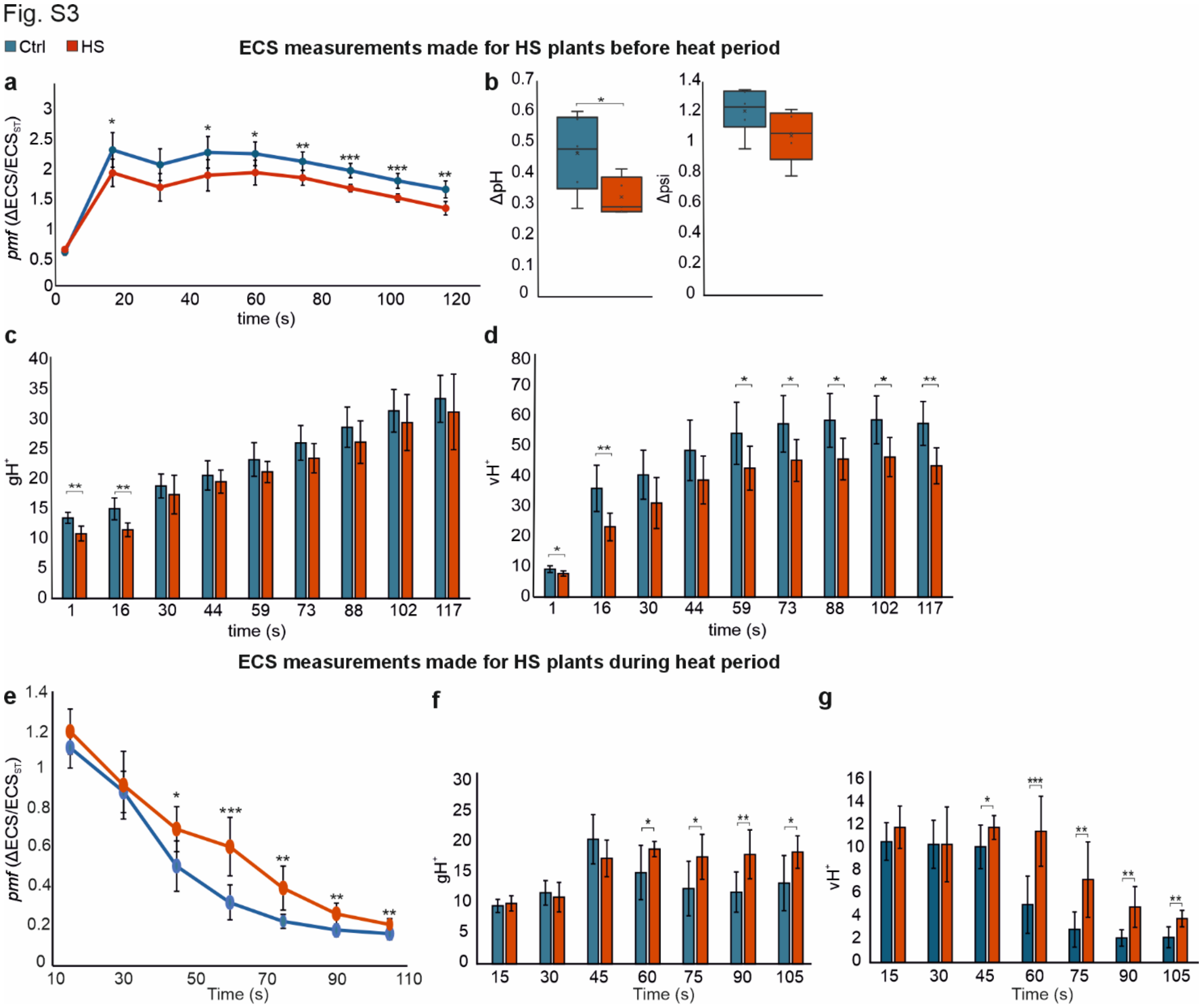


**Supplementary Figure 3.** ***pmf*** **was measured before the heat period (a – d) and in the middle of heat period at 38°C (e – g).** ECS was measured during transitions from dark to 130 µmol photons $m^{-2}$ $s^{-1}$ of red actinic light. Absorbance changes at 546 and 520 nm (electrochromic shift, ECS) was measured with a JTS-150 spectrometer. (a, e) Light-induced *pmf* was determined from the dark interval relaxation kinetics of the ECS signal. (b) Partitioning of *pmf* to ΔpH and ΔΨ after 2 min of illumination with 130 µmol photons $m^{-2}$ $s^{-1}$ actinic light. (c, f) Conductivity of the thylakoid membrane ($gH^+$) was determined as the inverse of the time constant of first-order post-illumination decay kinetics of the ECS signal. (d, g) Thylakoid proton flux ($vH^+$), calculated as *pmf* × $gH^+$. ECS measurements were done both before the heat period (a – d) and during the heat period (e – g) to distinguish acute short-term effects of heat treatment from long-term acclimatory responses. Statistical significance according to Student's t-test is indicated with an asterisk (* = < 0.05, ** = < 0.01, *** = <0.001). (n = 6 individual plants). Standard deviations are indicated with error bars. Ctrl = control plants, HS = heat-acclimated plants. Measurements were done at mid-day in the middle of heat period.

Fig. S4

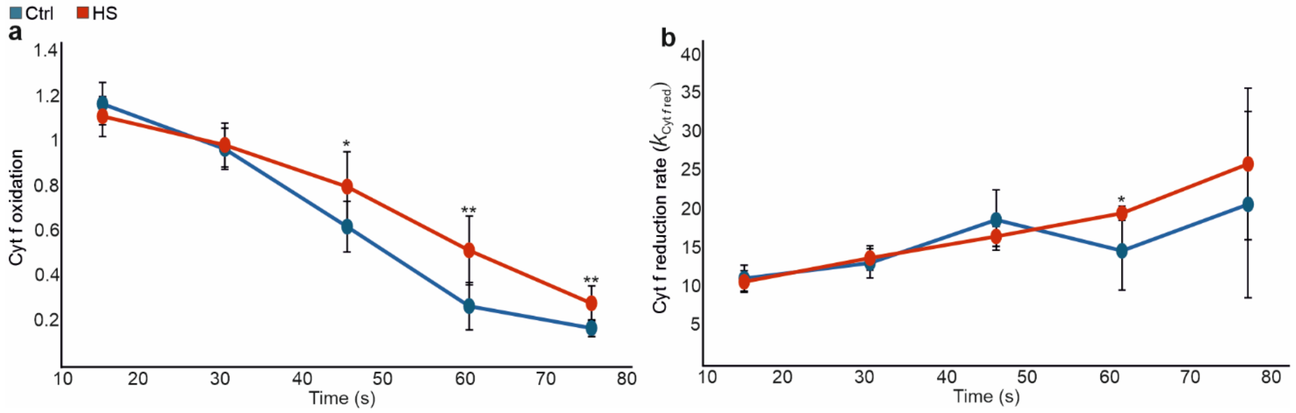


**Supplementary Figure 4**. a) Amount of oxidized cytochrome *f* during transitions from dark to 130 µmol photons $m^{-2}$ $s^{-1}$ of red actinic light. Cyt f redox changes were determined by monitoring absorbance changes at 554 nm with a baseline drawn between 546 and 573 nm. Amount of Cyt $f_{OX}$ was determined from the magnitude of the signal increase during 340 ms dark intervals, normalized by the initial decrease (oxidation) of the signal at the onset of illumination. b) Rate of cytochrome f reduction ($k_{Cyt\,f\,red}$) during transitions from dark to 130 µmol photons $m^{-2}$ $s^{-1}$ of red actinic light, determined from time constants of first-exponential fits the Cyt *f* signal kinetics during dark intervals. Statistical significance according to Student's t-test is indicated with an asterisk (* = < 0.05, ** = < 0.01, *** = <0.001). (n = 6 individual plants). Standard deviations are indicated with error bars. Ctrl = control plants, HS = heat-acclimated plants. Measurements were done at mid-day in the middle of heat period.

Fig. S5

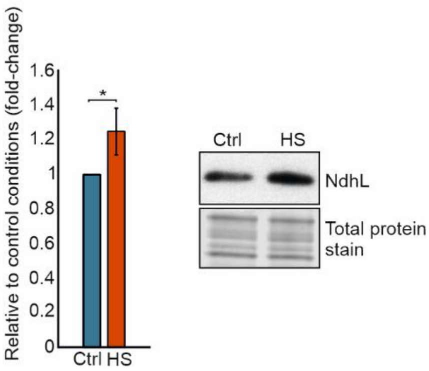


**Supplementary Figure 5.** Western blot and protein quantification from thylakoid membrane proteins from control (Ctrl) and heat-acclimated (HS) plants using antibody against NdhL. Thylakoid membrane proteins were isolated at the middle of heat period. Statistical significance according to Student's t-test is indicated with an asterisk (* = < 0.05). (n = 3 biological replicates). Standard deviations are indicated with error bars.

Fig. S6

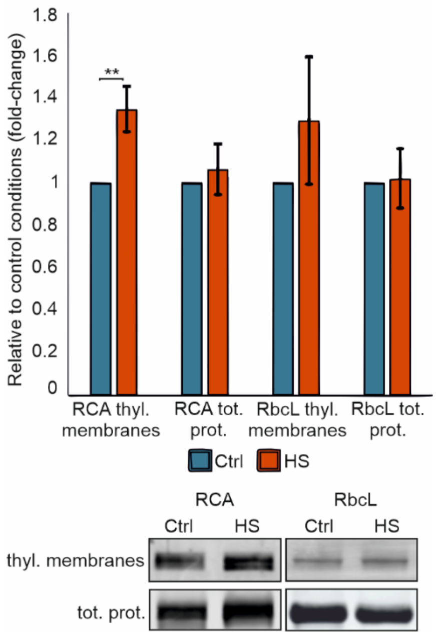


**Supplementary Figure 6.** Western blots and protein quantifications from thylakoids and total proteins of control (Ctrl) and heat-acclimated (HS) plants using antibodies against Rubisco large subunit (RbcL) and Rubisco activase (RCA). Thylakoid membrane proteins were isolated at the middle of heat period. Statistical significance according to Student's t-test is indicated with an asterisk (* =< 0.05, ** =< 0.01). (n = 3 biological replicates). Standard deviations are indicated with error bars.
